# Female reed warblers in social pairs with low MHC dissimilarity achieve higher MHC compatibility through random extra-pair matings

**DOI:** 10.1101/2023.04.17.537178

**Authors:** Lucyna Halupka, Emily O’Connor, Maria Strandh, Hanna Sztwiertnia, Ewelina Klimczuk, Dennis Hasselquist, Helena Westerdahl

## Abstract

Major Histocompatibility Complex (MHC) polymorphism is maintained by balancing selection through host-pathogen interactions and mate choice. MHC-based mate choice has been proposed across a wide range of vertebrates. However, the likelihood of its existence in songbirds has been questioned because of their poorly developed olfactory sense, which is a trait considered crucial in pre-copulatory mate choice to determine both own MHC and the MHC of putative partners. In this study, we show that female reed warblers, *Acrocephalus scirpaceus*, with extra-pair young in their nests have a lower MHC class I (MHC-I) dissimilarity with their social mate than females without extra-pair young in their nests. We also show that the MHC-I dissimilarity of successfully siring extra-pair males is not different from that of either the other males with territories surrounding the social nest (*i.e.* putative extra-pair males) or the pairs without extra-pair young in their nests. Taken together with the observation that extra-pair mating in reed warblers is common, we argue that these results support a scenario where extra-pair mating is more likely to lead to successful fertilisation when there is a high similarity in MHC-I between the female and her social male. Furthermore, as our data suggest that extra-pair mating at random can result in a higher MHC-I dissimilarity this scenario does not require any active female mate choice for MHC-I dissimilar males to drive this pattern.

## Introduction

The potential role of the Major Histocompatibility Complex (MHC) in mate choice has intrigued evolutionary biologists and behavioural ecologists for decades (Geßner, Nakagawa, Zavodna, & Gemmell, 2017; Kamiya, O’Dwyer, Westerdahl, Senior, & Nakagawa, 2014; Milinski, 2006; Penn & Potts, 1999). MHC molecules are central for antigen presentation and T-cell mediated adaptive immune responses (Neefjes, Jongsma, Paul, & Bakke, 2011). MHC genes are highly variable and this polymorphism is maintained by selection from pathogens and / or mate choice (Tobias L. Lenz, 2011; Penn & Potts, 1999; Piertney & Oliver, 2006). An optimally high MHC diversity (the number of different MHC alleles per individual), or divergence (functional distance between MHC alleles within an individual), is advantageous since a wider range of pathogens can be detected and eliminated, and studies have shown that MHC diversity / divergence correlate with, disease resistance, increased survival and fitness in wild populations (T. L. Lenz, Mueller, Trillmich, & Wolf, 2013; Radwan et al., 2012; Roved, Hansson, Tarka, Hasselquist, & Westerdahl, 2018; Wakeland et al., 1990). To maximize the MHC diversity of offspring females should chose partners with the highest MHC diversity whereas to maximize the MHC divergence of offspring females should select males with the most dissimilar MHC genotype relative to her own, *i.e*., disassortative MHC-based mating (T. L. Lenz et al., 2013; Pineaux et al., 2022; Roved et al., 2018; Wakeland et al., 1990). Note, that several studies have also tested for a mate choice to gain an optimal rather than a maximal MHC diversity / MHC dissimilarity with partners (e.g. Rekdal, Anmarkrud, Lifjeld, & Johnsen, 2019).

A large number of studies have investigated whether social mate choice is MHC-based (MHC dissimilarity, MHC diversity and specific MHC genotypes), using a plethora of study systems, techniques and approaches (Bererhi et al., 2023; Bollmer, Dunn, Freeman-gallant, & Whittingham, 2012; Gasparini, Clelia, Congiu, Leonardo, Pilastro, 2015; Geßner et al., 2017; Milinski, 2006; Potts, Wayne K. Manning, C. Jo. Wakeland, 1991; Strandh et al., 2012; Helena Westerdahl, 2004). In general, MHC-based social mate choice appears to be somewhat infrequent among vertebrates (Chaix, Cao, & Donnelly, 2008; Derti, Cenik, Kraft, & Roth, 2010; Kamiya et al., 2014; Mays, Albrecht, Liu, & Hill, 2008). However, social mate choice in many species is strongly influenced by direct benefits, such as territory quality, social rank, and ability to provision for offspring, which may override or obscure the signal of MHC-based mate choice, an indirect benefit (Duffy, Ball, Duffy, & Ball, 2002; Griffith, Owens, & Thuman, 2002; Hasselquist, Dennis, Staffan Bensch, 1996; Hasselquist, 1998). Extra-pair mate choice in contrast, *i.e.,* mating outside the social pair-bond, is largely free of such constraints and therefore provides an excellent model for investigating MHC-based mate choice (Kamiya et al., 2014; Zelano & Edwards, 2002). Indeed, extra-pair mate choice for either MHC diversity or MHC dissimilarity could partly compensate for low MHC diversity in the social male or low MHC dissimilarity within the social pair.

Songbirds are well suited to studies of extra-pair mate choice, as although most species are socially monogamous, extra-pair mating is common (Davies, Butchart, Burke, Chaline, & Stewart, 2003; Halupka, Sztwiertnia, Borowiec, Klimczuk, & Leisler, 2014; Hasselquist & Sherman, 2001). Many songbird species have highly duplicated MHC genes, which has previously made obtaining accurate MHC genotypes challenging (Davies et al., 2003; Halupka et al., 2014). Since the advent of next-generation sequencing, the use of amplicon high-throughput sequencing (HTS) has enabled the detailed characterization of MHC genes even in species with extreme MHC gene copy numbers (Halupka et al., 2021; Winternitz et al., 2015). A handful of studies have used HTS approaches to study the role of MHC in mate choice in songbirds, though with mixed results (for exampe Bollmer et al., 2012; Rekdal et al., 2019; Winternitz et al., 2015). Therefore, many open questions surrounding the role of MHC genes in mate choice in songbirds remain, such as whether extra-pair mating compensates for suboptimal social mate choice in relation to MHC dissimilarity (i.e., the functional distance based on the peptide-binding regions of all MHC alleles within a pair).

Here we test whether MHC class I (MHC-I) dissimilarity plays a role in social and extra-pair mate choice in a wild study population of reed warblers, *Acrocephalus scirpaceus,* in South-West Poland. Reed warblers are socially monogamous but have a high frequency of extra-pair mating (up to 33% of breeding females have at least one extra-pair chick in their brood) (Halupka et al., 2014). We combined field observations of breeding pairs over six years (128 females and 127 males) with data on their MHC-I genotypes to explore the relationship between MHC-I dissimilarity within social pairs, the occurrence of extra-pair paternity and the choice of extra-pair male. This extensive dataset allowed us to demonstrate how the choice of extra-pair male relates to the locally available males in terms of MHC-I dissimilarity. In doing so, we present compelling evidence for how extra pair mating can compensate for poor within-pair MHC-I dissimilarity in the absence of active extra-pair mate choice.

## Materials and Methods

### Study population

Reed warblers are socially monogamous migratory passerine birds (Halupka et al., 2021). Most social pairs remain together throughout a breeding season (May – August) laying up to four clutches, but disband between seasons (Halupka et al., 2014). Both males and females are heavily engaged in both incubation and feeding young (Klimczuk et al., 2015). Each female generally has only a few unmated males to choose from when seeking a social mate. However, there are often many male neighbours in the vicinity of any given female’s nest, providing opportunities for extra-pair mating. During May to August of 2006 to 2011, we studied a population of reed warblers breeding in the nature reserve "Stawy Milickie”, SW Poland (51°32 N, 17°20 E). We monitored the study plot daily from the time the first individuals arrived and mist-netted and individually marked birds throughout the breeding season. Each year between 29 and 51 breeding pairs were monitored, as well as putative extra-pair males around each nest.

### DNA extraction

A total of 288 adults were sexed and a blood sample (10*–*40 µl) was taken from each. Nestlings were blood sampled when 6-8 days old (501 chicks). Blood samples were preserved in 96% ethanol and kept in a freezer until DNA extraction using DNeasy Blood and Tissue Kit (QIAGEN, Hilden, Germany) or GeneMATRIX Tissue DNA Purication Kit (EURx, Gdańsk, Poland). The DNA extracts were kept frozen at -20 °C until used (QIAGEN, Hilden, Germany).

### MHC-I genotyping

We prepared forward and reverse 6bp-tagged amplicon libraries for high-throughput 454 sequencing of exon 3, the most variable fragment of the gene encoding the antigen binding site of the MHC-I molecule, from 288 adult individuals (24 individuals were duplicated to aid with filtering the data, n = 312) using the forward primer HN36 and reverse primer HN46 (the amplicon without primers is 257 bp and the complete exon 3 is 274 bp (H. Westerdahl, Wittzell, Von Schantz, & Bensch, 2004; H Westerdahl, Wittzell, & von Schantz, 1999)). These primers were previously designed to amplify MHC-I alleles in a closely related species, the great reed warbler, *Acrocephalus arundinaceus*, and we ran PCRs as described in (Roved et al., 2018). The 454-sequencing data (312 amplicons, 46471 reads) was filtered to remove low-quality amplicons and sequences, *i.e.*, sequences representing artefactual MHC-I alleles and potentially non-functional alleles as described in (Roved et al., 2018). The minimum total sequence abundance/individual (filter 1) was set to 75 reads and the minimum abundance of one sequence in all individuals (filter 2) was set to 4 reads. The 24 technical duplicates were used to determine the threshold for minimum relative abundance of a sequence/individual (filter 3). This filter was set to 6% (based on best between-duplicate identity: 95% total similarity). Any alleles not found in at least two independent PCR reactions were removed. Sequences were inspected manually and any non-functional sequences (four) or chimeras (two) were removed. After filtering, the final data set contained 281 sequences found in 255 individuals (128 females and 127 males) among which there were 154 known breeding pairs (12 to 36 pairs per year). The average number of reads per amplicon was 109 and each individual had 1-9 MHC-I exon 3 alleles.

Of the 281 nucleotides sequences, six had 100% identity and at least 200 base pair overlap with MHC-I exon 3 sequences found previously in reed warblers (GenBank accession numbers: KU169387, KU169375, KU169386, KU169388, KU169376 and KU169393). The sequences of the remaining 275 nucleotide sequences were uploaded to GenBank (accession numbers: OR053210 to OR053484).

### Parentage analysis

Parentage was analysed for 501 chicks. We used six fluorescently labelled microsatellites (Ase18, Ase25, Ase37, Ase48, Ase 58, Ppi2) in parents and offspring of 154 pairs, following methods in (Davies et al., 2003). The PCR products were fragment analysed on a DNA sequencer ABI PRISM® 3100-AvantTM Genetic Analyser (Applied Biosystems) and the resulting chromatograms were analysed using ABI Genescan software. Exclusion probabilities were calculated with Cervus 3.0, and paternity analysis was conducted using the GeneMapper software (version 4.0). Extra-pair young were found in 8.8 - 34.3 % of the nests across years. In total 35 pairs had extra-pair young in their nest: 21 extra-pair males were identified (across 18 social pairs) and for 17 social pairs the extra-pair male could not be identified. There were 106 pairs with no extra-pair young in their nest.

### Calculating functional MHC-I distances as a measure of dissimilarity in pairs

In order to assess MHC dissimilarity (MHC-I functional distance) of all known reed warbler pairs we estimated MHC-I functional distances according to (Strandh et al., 2012). We computed the MHC-I functional distances as follows: The verified MHC-I sequences (N = 281) were translated into a set of 243 unique amino acid sequences and from these we extracted the MHC-I exon 3 peptide-binding regions (17 amino acid PBR positions, inferred from (Bjorkman et al., 1987)) that made up a set of 160 unique PBR amino acid sequences. The PBR sequences were represented by five physicochemical descriptors per amino acid (Sandberg, Eriksson, & Sjo, 1998), that were subsequently used to construct a maximum likelihood tree with contml in the PHYLIP package, v 3.69. We used this tree (Fig. S1), as a reference from which the overall MHC-I functional distances between pairs (considering the full MHC-I repertoire of each male and female) were extracted with the UniFrac software (Lozupone & Knight, 2005). MHC-I functional distances were compiled for:; (1) *social pairs with extra-pair young*: **SP-EPY** (N = 35 pairs), (2) *extra genetic pairs*: **EGP**, the corresponding extra-pair dyads, when known, to the SP-EPY social pairs (N=21 dyads corresponding to 18 of the SP-EPY as a result of some males being the EGP for more than one SP-EPY), (3) *potentially available extra-pair males* (**PA-EPM**) with a territory within a 60 m radius of SP-EPY nests where the EGP was known, as all EGP males had a territory within 60 m of the SP-EPY nest (6–14 putative males per SP-EPY) and (4) *social pairs with only within-pair young:* **SP-WPY,** i.e., pairs without EPY in the nest (N = 106 pairs). In total, MHC-I genotypes were available for 128 females and 127 males in the population enabling MHC-I functional distances to be calculated between 16256 theoretically possible pairs.

### Statistical analyses

To test whether there was evidence of non-random pairing regarding MHC-I dissimilarity for the different pair types (SP-WPY, SP-EPY of EGP) the position of the mean MHC-I dissimilarity for each of the observed groups was compared against the distributions of mean MHC-I dissimilarity from 10000 groups of random pairs (matched for the group size of the observed means i.e.,106 for SP-WPY, 35 for SP-EPY and 21 for EGP). Random pairs were sampled from all possible combinations of 128 females and 127 males genotyped in the population (i.e., 16256 theoretically possible pairs) with replacement between (not within) each of the 10000 iterations for each group size, to ensure a broad sampling of all possible pairs. If the observed mean MHC-I dissimilarity fell outside the two-tailed 95% confidence intervals of the random distribution, then the observed MHC-I dissimilarity was significantly different to that expected from random pairing given the group size at the level of P < 0.05. Evidence of non-random pairing regarding MHC-I diversity was tested in an identical fashion but using the number of MHC-I alleles of males instead of MHC-I dissimilarity within pairs. To test whether females in SP-EPY chose extra-pair males in a non-random fashion regarding MHC-I dissimilarity, we generated 10000 random pairings between each female and all the surrounding males, i.e., PA-EPM with a territory within 60 m of the nest. We then tested whether the MHC-I dissimilarity of the observed EGP fell outside the two-tailed 95% confidence intervals of the distribution of the random pairings. The randomisation analyses were conducted in R using base code (*see R scripts in supplementary files*).

Differences in the MHC-I dissimilarity scores between pair types (SP-WPY, SP-EPY of EGP) were analysed with general linear mixed models (GLMM) using the R package lme4 version 1.1-29 (Bates, Mächler, Bolker, & Walker, 2015). The response variable was MHC-I dissimilarity scores, with pair type as a three-level fixed effect. Female identity and year were included as random effects in the models to account for the non-independence of pairings within the same year and the same female with different male partners. To test whether MHC-I dissimilarity differed between SP-EPY and the EGP, the same model was run on a subset of the data that only included the paired results from SP-EPY where the EGP was known (18 SP-EPY and 21 EGP). P-values to indicate the significance of fixed effects in the GLMMs were obtained using parametric bootstrapping (1000 simulations) implemented in the R package afex (Singmann H, Bolker B, Westfall J, Aust F, 2023). To ensure that any observed patterns in MHC-I dissimilarity between pairs was not driven by a choice for males with high MHC-I diversity we tested for differences in the number of MHC-I alleles of males across the different pair types. These analyses were performed in a similar fashion using generalised linear mixed models in lme4 and specifying a Poisson distribution to account for count data. We also checked whether there was an association between MHC-I dissimilarity and the number of MHC-I alleles of males within pairs using a Spearman’s rho rank correlation test performed in the R package Hmisc V. 5.0-1 (Harrell F. Jr, 2023).

To examine the MHC-I dissimilarity of the SP-EPY and the EGP in the context of the putatively available extra-pair males (PA-EPM), we used the MHC-I genotypes of the surrounding males. MHC-I dissimilarity scores were calculated between each focal female and all her putatively available extra-pair males (all males with a territory within a 60 m radius of the nest). This data was then used to calculate the percentile of the MHC-I dissimilarity of the SP-EPY and EGP within the distribution of the MHC-I dissimilarity offered by the surrounding males. For example, if the percentile of the SP-EPY was 0.2, then the MHC-I dissimilarity of the SP-EPY was lower than 80% of the potential pairings between the focal female and the other surrounding males. The difference between the percentile of the observed pairs MHC-I dissimilarity scores and that of the median (*i.e.*, 0.5) was tested using GLMMs. The response variable was MHC-I dissimilarity with the observed percentile (either SP-EPY or EGP) or median treated as a two-level fixed factor and pair included as a random effect.

All analyses were conducted in R version 4.1.1 (R Core Team, 2023)

## Results

MHC-I dissimilarity (functional distance) was calculated for three different pair types: social pairs with extra-pair young (**SP-EPY**, N = 35 pairs); extra genetic pairs (**EGP**, N=21 occurrences) and social pairs with only within-pair young (**SP-WPY**, i.e. pairs without EPY in the nest, N = 106 pairs). The mean MHC-I dissimilarity within each of the three pair types was tested against what would be expected from random mating and the only group with MHC-I dissimilarity scores outside what would be expected from random mating were the SP-EPY, which had a significantly lower mean MHC-I dissimilarity score than would be expected by chance from 35 pairs (Fig. 1).

**Figure 1:**
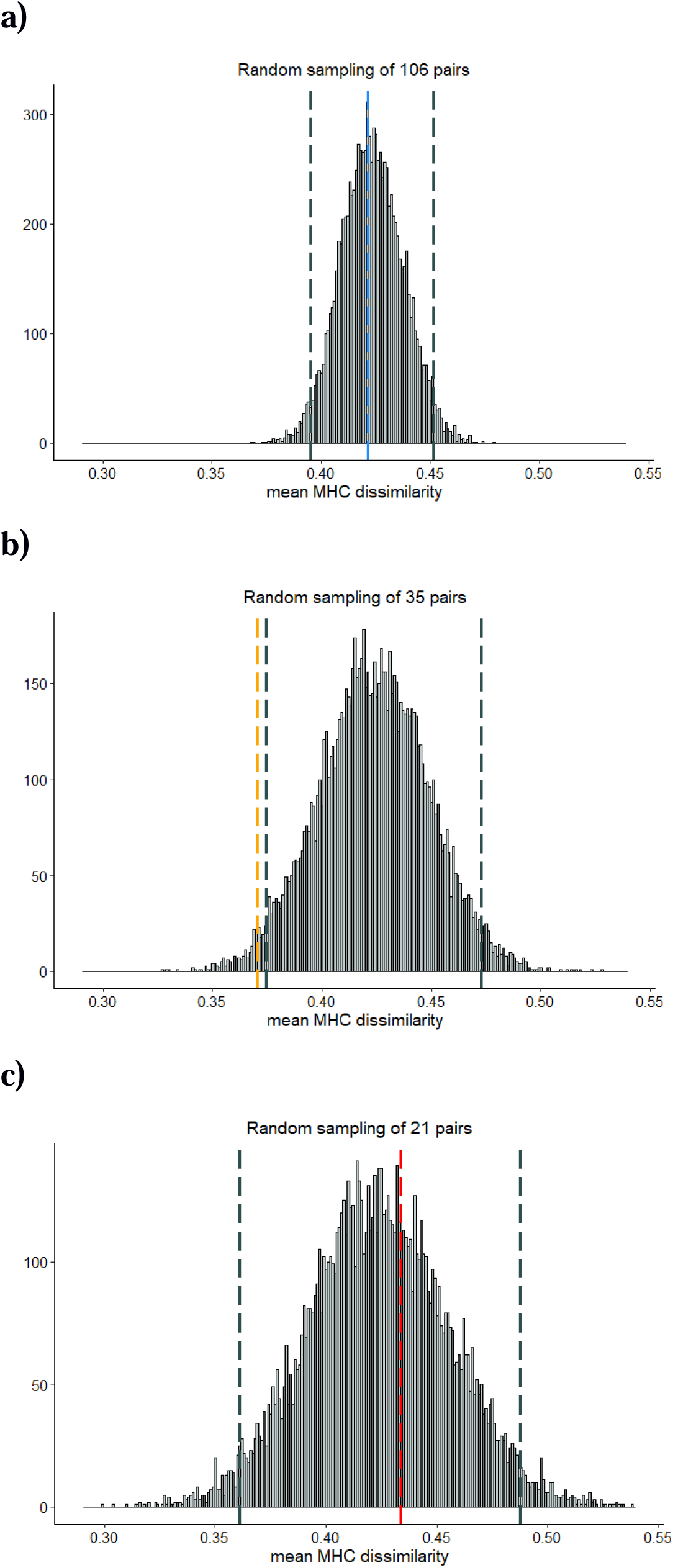
Distribution of 10000 mean MHC-I dissimilarity scores calculated from randomly selected groups of pairs using all MHC-I genotyped individuals in the population (128 females and 127 males). Group sizes were matched for the sample size of the comparison dataset: (**a**) 106 pairs for the social pairs with only within-pair young SP-WPY, (**b**) 35 pairs for the social pairs with extra-pair young SP-EPY and (**c**) 21 pairs for extra genetic pairs EGP. Dark grey dashed lines depict the lower 2.5% and upper 97.5% of the data in both figures (95% confidence intervals). The mean MHC-I dissimilarity for the SP-WPY, SP-EPY and EGP are depicted by the blue, orange and red dashed lines, respectively.

The randomisation test suggests that SP-EPY are more similar than expected under random mating. However, there was no overall significant difference in the MHC-I dissimilarity of the three different pair types (SP-EPY, SP-EPY and EGP, Fig. 2, χ^2^ _2, 156_ = 3.75, P = 0.15). Though, there was a trend for SP-EPY to have lower MHC-I dissimilarity than the corresponding EGP in the cases where the EGP could be identified (Fig. 2, χ^2^ _1, 37_ = 3.09, P = 0.08). Indeed, after removing an outlying data point with extremely high MHC-I dissimilarity from SP-EPY and its corresponding EGP, this difference became statistically significant (χ^2^ _1, 35_ = 6.04, P = 0.01). There was no significant difference in MHC-I diversity (the number of different MHC-I alleles in males) between any of the pair-types (Fig S2, SP-EPY, SP-EPY and EGP: χ^2^ _2, 157_ = 1.36, P = 0.51; SP-EPY vs known EGP: χ^2^ _1, 38_ = 1.65, P = 0.20), and no overall correlation between MHC-I diversity of males and MHC-I dissimilarity within pairs (r = 0.05, P = 0.51, n = 188), showing that male MHC-I diversity does not explain MHC-I dissimilarity in pairs. Furthermore, there was no evidence of non-random mate choice with respect to the number of MHC-I alleles of males (Fig. S3).

**Figure 2:**
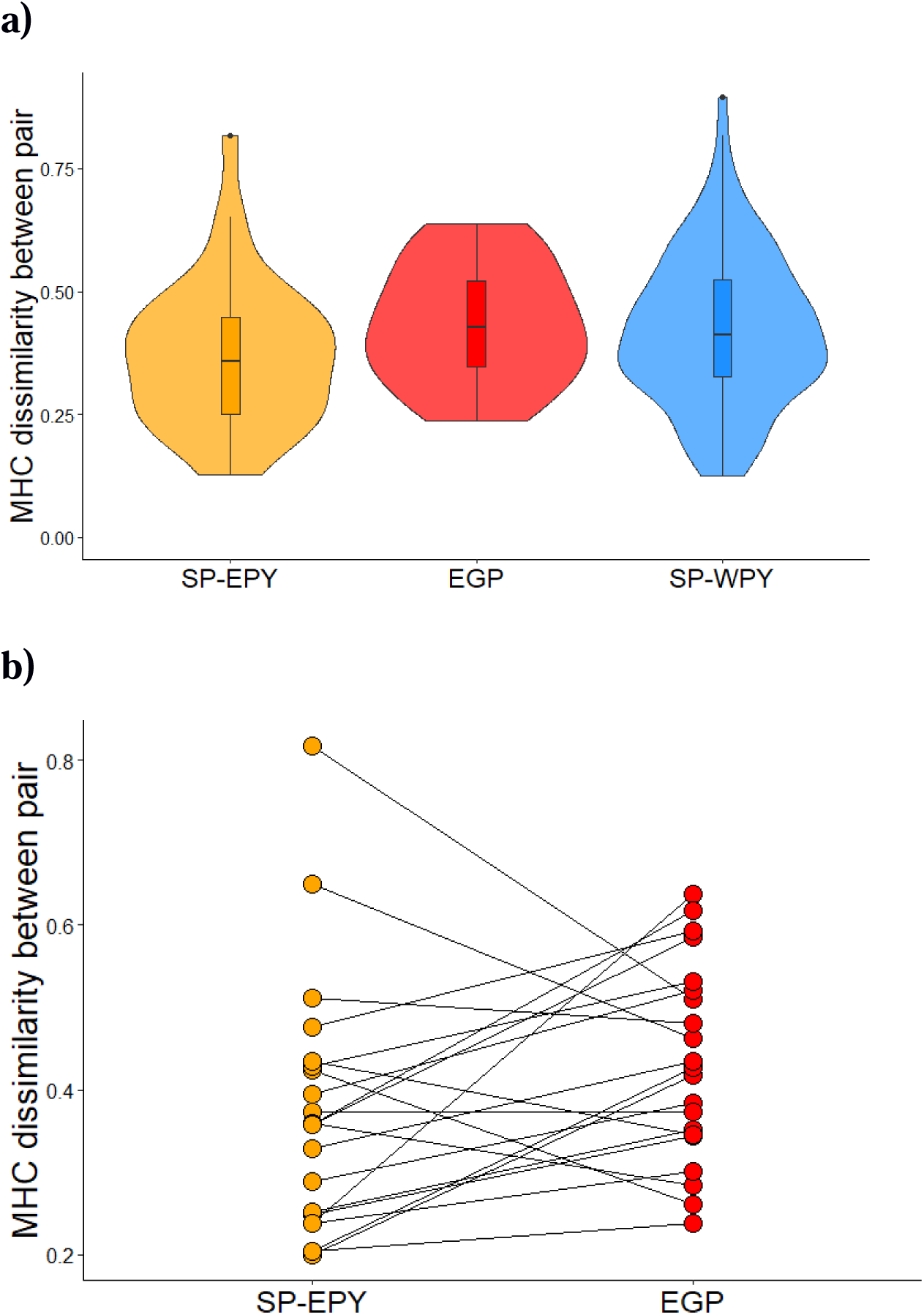
MHC dissimilarity scores between different pair types. **a)** Violin plot of the MHC dissimilarity between social pairs with extra-pair young (SP-EPY, n = 35), females with extra-pair young and their known extra-pair male (extra genetic pair, EGP, n = 21) and social pairs with only within-pair young in their nest (SP-WPY, n = 106). Boxplots within the violins (central line = median, whiskers = 1.5 times the interquartile range of the data, black dots = outliers). **b)** Comparison of the MHC dissimilarity between SP-EPY where the extra-pair male is known (n = 18) and known EGPs (n = 21). Lines link the female with the corresponding extra-pair males.

Both the randomisation tests and the analyses comparing the pairwise MHC-I dissimilarity of different pair types suggest that SP-EPY have lower MHC-I dissimilarity than other pair types. To analyse this finding further, we compared the MHC-I dissimilarity of SP-EPY and the EGP with all available extra-pair males within a 60 m radius of SP-EPY nests where the EGP was known. The MHC-I dissimilarity of SP-EPY was significantly lower than the median MHC-I dissimilarity of the female if paired with any of the other surrounding males (Fig. 3A, χ^2^ _1, 38_ = 5.04, P = 0.02), whereas no such difference was evident between EGP and the female if paired with any of the other surrounding males (Fig. 3B, χ^2^ _1, 38_ = 2.03, P = 0.15). Furthermore, there was no evidence that females in SP-EPY selected the males in their EGP in a non-random fashion, given the potential MHC-I dissimilarity represented by the surrounding males (Fig. S4).

**Figure 3:**
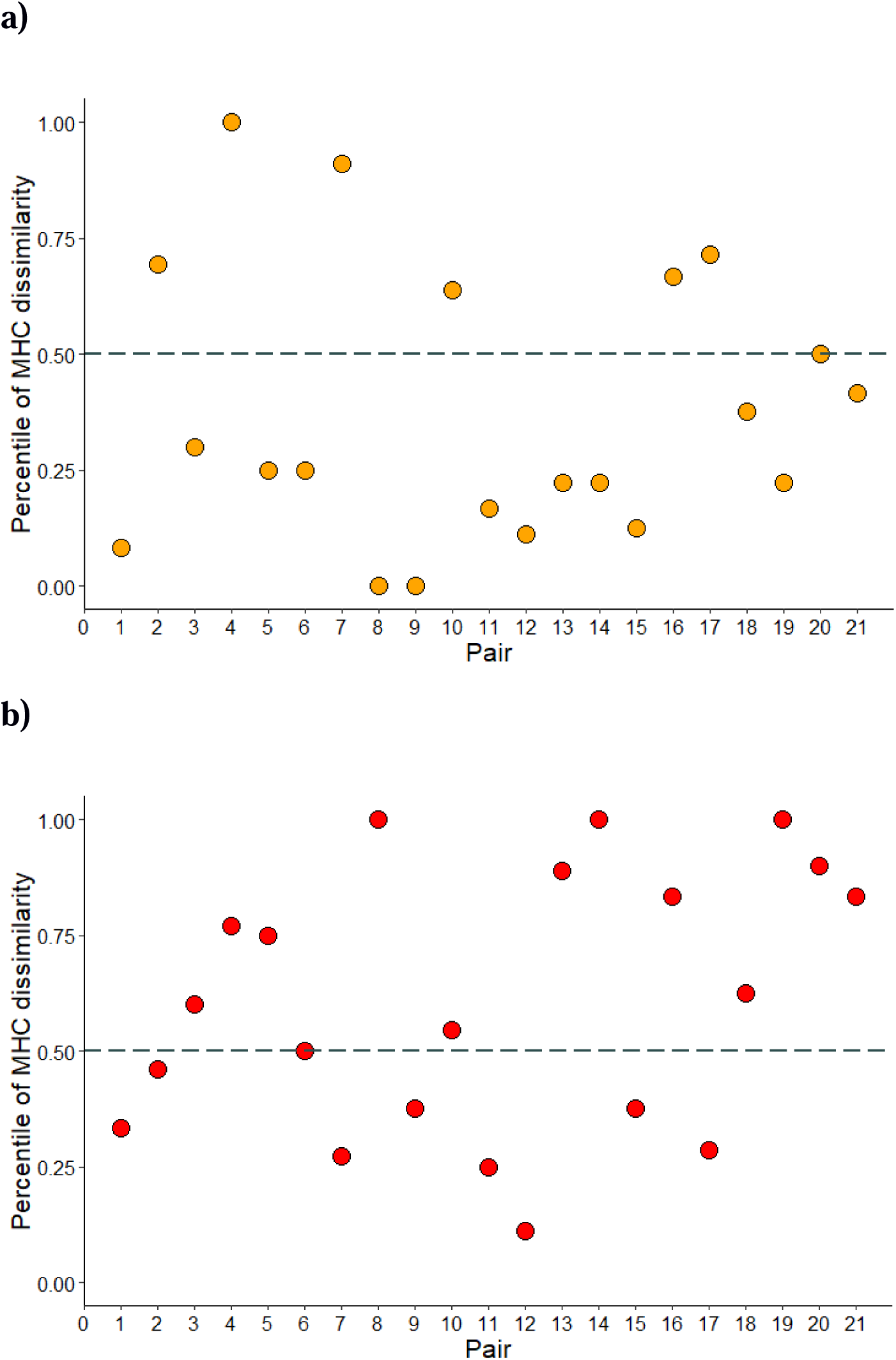
The percentile of the MHC dissimilarity of (**a**) social pairs with extra-pair young SP-EPY and (**b**) extra genetic pairs EGP within the distribution of MHC dissimilarity between the focal female and all potentially available extra-pair males within a 60m radius of her nest. Dark grey dashed line denotes the median.

## Discussion

Overall, our analyses suggest that female reed warblers in social pairs with low MHC-I dissimilarity have more successful extra-pair matings than females in social pairs with higher MHC-I dissimilarity. However, these extra-pair matings seem to occur at random with respect to MHC-I dissimilarity among the putative extra-pair males available close to the social nest.

Reed warbler pairs with extra-pair young in their clutches (social-pairs with extra-pair young, SP-EPY) had significantly lower MHC-I-dissimilarity than expected from random pairing. Furthermore, MHC-I dissimilarity tended to be higher between females and their extra-pair males (extra-genetic pairs, EGP) than in SP-EPY, hence females in pairs with low MHC-I dissimilarity achieve improved MHC-I compatibility through extra-pair mating. Therefore, low MHC-I dissimilarity within social pairs is associated with paternity loss for the social males in this population of reed warblers. We found no evidence of any difference in MHC-I diversity (the number of MHC-I alleles per individual in males) between SP-EPY and EGP. This is important as it demonstrates that the difference we observed in MHC-I dissimilarity between SP-EPY and EGP is not driven by males in EGP having higher MHC-I diversity, which has been shown in the closely related Seychelles warbler, *Acrocephalus sechellensis*, as well as the scarlet rose finch, *Carpodacus erythrinus* (Brouwer et al., 2010; Promerová et al., 2011; Richardson, Komdeur, Burke, & von Schantz, 2005). We also found no evidence of non-random pairings regarding the number of MHC-I alleles in males, further suggesting that MHC-I diversity does not play an important role in mate choice in this population of reed warblers.

The high frequency of nests with EPY in reed warblers (up to 33%) indicates that extra-pair mating is common in this population (Halupka et al., 2014). If most females engage in extra-pair mating, then these copulations appear to result in extra-pair offspring more frequently when the MHC-I dissimilarity of the social pair is low. In such a scenario, post-copulatory mechanisms (*e.g.,* cryptic female choice or sperm competition) could select for successful fertilisation by the extra-pair male as a result of higher MHC-I compatibility between the female and the extra-pair male compared to the social male (Tregenza & Wedell, 2000). There is evidence for post-copulatory MHC-I-based mate choice in birds: an experimental study in domestic chickens, *Gallus gallus domesticus*, showed that MHC-I dissimilarity predicted fertilisation success (Løvlie, Gillingham, Worley, Pizzari, & Richardson, 2013). Moreover, MHC dissimilarity has been shown to influence fertilisation success in Chinook salmon *Oncorhynchus tshawytscha* (Geßner et al., 2017), mice (Potts, Wayne K. Manning, C. Jo. Wakeland, 1991; Wedekind, Chapuisatt, Macas, & Rulicke, 1996) and even humans (Jokiniemi, Kuusipalo, et al., 2020; Jokiniemi, Magris, et al., 2020). The observation in this study, that the EGP and social-pairs with only within-pair young (SP-WPY), had similar MHC-I dissimilarity whereas SP-EPY tended to have lower MHC-I dissimilarity supports the concept that MHC-I compatibility may also play a role in fertilization success in reed warblers.

Only the SP-EPY showed evidence of non-random mating in respect to MHC-I, with lower-than-expected MHC-I dissimilarity. This is likely to reflect the constraints associated with social pair formation, such that a larger proportion of females will be paired with suboptimal males in socially monogamous species (Hasselquist & Sherman, 2001). Overall, there was little evidence to support any active mate choice for MHC-I dissimilar males in this study. Although the EGP had higher MHC-I dissimilarity than the SP-EPY, there was no indication that the female chose an extra-pair male with particularly high MHC-I dissimilarity given the surrounding putatively available males. The MHC-I dissimilarity of the EGP was not in the upper range of the MHC-I dissimilarity that could potentially have been achieved by the female if she copulated with a different locally available male. Furthermore, there was no evidence of non-random pairing with the EGP with respect to the MHC-I dissimilarity of surrounding males. A lack of active choice for MHC-I dissimilar males when engaging in extra-pair copulations is consistent with our observation that extra-pair mating is extremely time-constrained in reed warblers, since the social male mate-guards the female (Halupka unpublished data). Extra-pair mating may therefore be opportunistic and lack discerning choice. Indeed, given that the MHC-I dissimilarity of the SP-EPY was significantly below the median MHC-I dissimilarity of the focal female with other surrounding males, it is likely that most locally available males represent an improved match in terms of MHC-I dissimilarity compared to the social male. Therefore, even extra-pair mating at random is likely to result in an improved match in terms of MHC-I dissimilarity.

We cannot exclude the possibility that the higher MHC-I dissimilarity of the EGP compared to SP-EPY reflects inbreeding avoidance rather than specific MHC-I effects. However, we consider this alternative explanation unlikely as the low natal fidelity of this population of reed warblers is likely to result in a highly outbred population (Halupka unpublished data).

Our results raise the intriguing possibility that reed warbler females who are in a social pair with poor MHC-I compatibility improve the MHC-I repertoire of her offspring through random extra-pair mating. Whilst previous work has suggested that extra-pair mating may be a mechanism by which the MHC-I repertoire of offspring is improved in passerine birds (Bollmer et al., 2012; Rekdal et al., 2019; Winternitz et al., 2015), here we provide evidence that this could be effectively achieved through random extra-pair mating. When MHC-I dissimilarity with the social mate is low, any other male may represent a better prospect for the female in terms of MHC-I compatibility.

## Acknowledgements

We are grateful to Giulia Casasole, Beata Czyż, Alicja Dziachan, Magdalena Soboń, Stanisław Rusiecki and Łukasz Tomasik for their help in the field, to Jörgen Ripa, Konrad Halupka and John Hutchinson for statistical support. This study was supported by grants from the Swedish Research Council (Vetenskapsrådet; grants no. 2016-04391 and 2020-03976 to DH, 2015-05149 and 2020-04285 to HW) and the European Research Council (ERC) under the European Union’s Horizon 2020 research and innovation program (ERC advanced grant no. 742646 to DH).

## Data Accessibility and Benefit-Sharing

All raw data used in this study and associated metadata is uploaded to a Dryad repository (XXXX). The sequences of the 281 MHC-I alleles can be found on GenBank (accession numbers: OR053210 to OR053484, KU169387, KU169375, KU169386, KU169388, KU169376 and KU169393).

## Author contributions

LH designed the project, performed the fieldwork and MHC lab analyses, prepared the data for the statistical analyses, and contributed to the interpretation of the data. EO performed the statistical analyses and contributed to the interpretation of the data. MS performed the functional analyses of the MHC data and contributed to the interpretation of the data. HS performed fieldwork and microsatellite analyses. EK performed the fieldwork. DH contributed to the interpretation of the data. HW helped to plan the project, supervised the project, contributed to the statistical analyses and to the interpretation of the data, all co-authors contributed to writing of the article.

## Notes

### Competing Interest Statement

The authors have declared no competing interest.

### Summary of Updates

The text in this version has been slightly updated and new references added.

