## Supplementary Material for "Female reed warblers in social pairs with low MHC dissimilarity achieve higher MHC compatibility through random extra-pair matings"

**Supplementary figures**

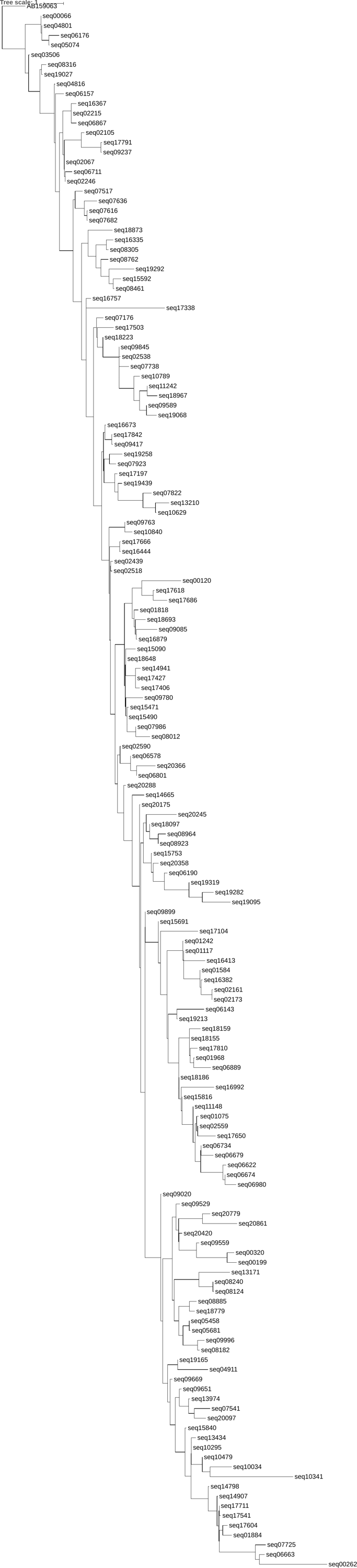

**Figure S1.** Maximum likelihood tree based on 160 functional MHC class I exon 3 sequences from the reed warbler and one MHC class I sequence from the chicken, Gallus gallus, (Acc nr AB159063) as an outgroup. The peptide binding region was first transformed to physiochemical z-descriptors used to construct the tree with the software contml (PHYLIP-package, v 3.69) using default settings for continous characters.

**a)**

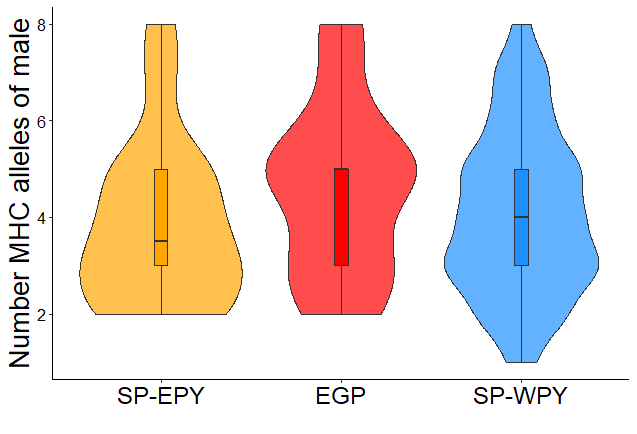

**b)**

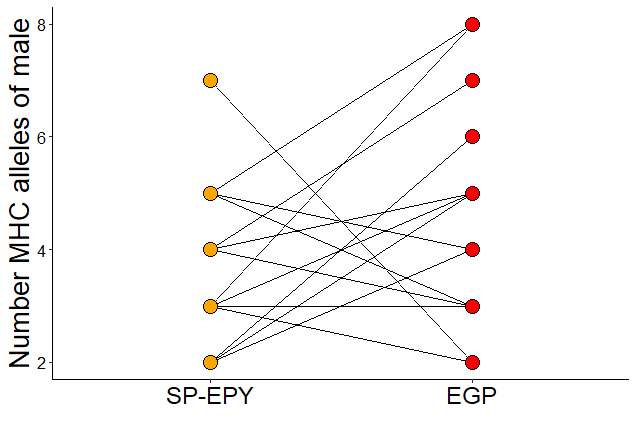

**Figure S2.** Number of MHC alleles of male within different pair types. a) Violin plot of the number of MHC alleles of males in social pairs with extra-pair young (SP-EPY, n = 35), females with extra-pair young and their known extra-pair male (extra genetic pair, EGP, n = 21) and social pairs with only within-pair young in their nest (SP-WPY, n = 106). Boxplots within the violins (central line = median, whiskers = 1.5 times the interquartile range of the data, black dots = outliers). b) Comparison of the number of MHC alleles of males in SP-EPY where the extra-pair male is known (n = 18) and the known EGP male (n = 21). Lines link the female with the corresponding extra-pair males.

**a)**

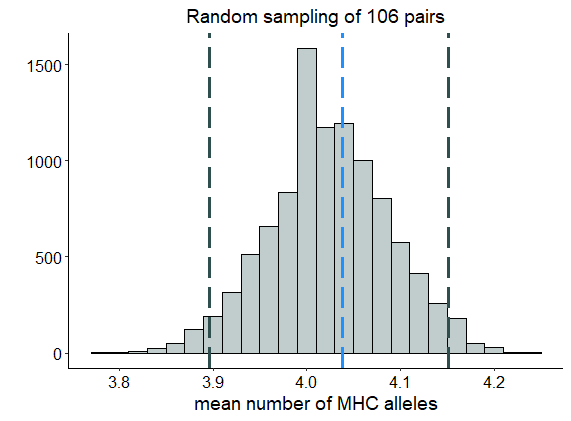

**b)**

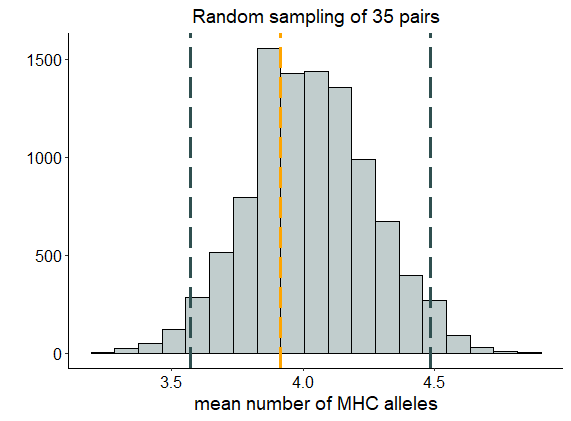

**c)**

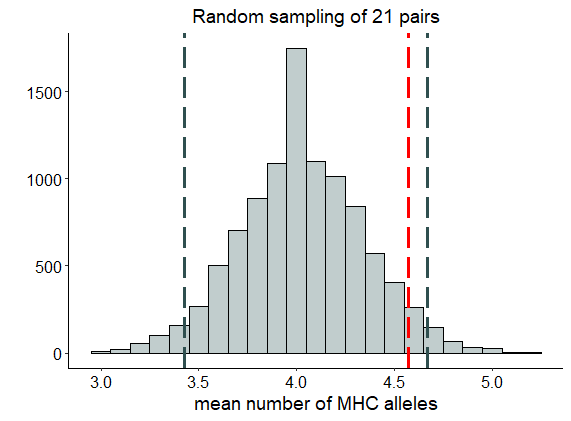

**Figure S3:** Distribution of 10000 mean number of MHC-I alleles per male calculated from randomly selected groups of males using all 127 MHC genotyped males in the population. Group sizes were matched for the sample size of the comparison dataset: (**a**) 106 males for the social pairs with only within-pair young SP-WPY, (**b**) 35 males for the social pairs with extra-pair young SP-EPY and (**c**) 21 males for extra genetic pairs EGP. Dark grey dashed lines depict the lower 2.5% and upper 97.5% of the data in both figures (95% confidence intervals). The mean number of MHC-I alleles in males for the SP-WPY, SP-EPY and EGP are depicted by the blue, orange and red dashed lines, respectively.

| a.   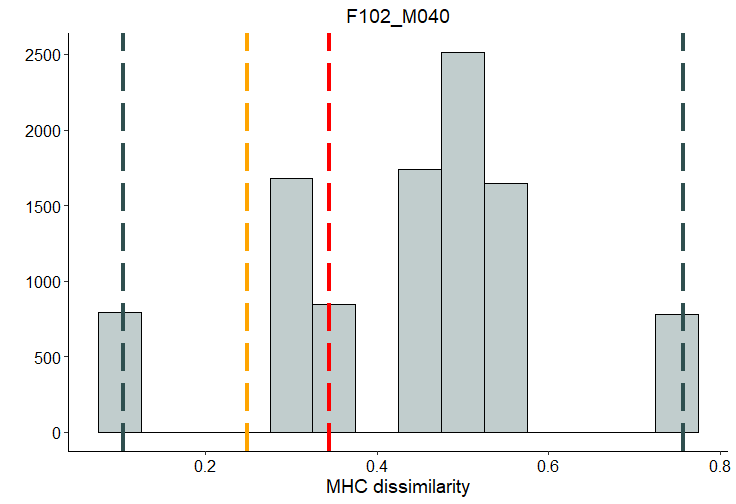  n = 12 | **b.**  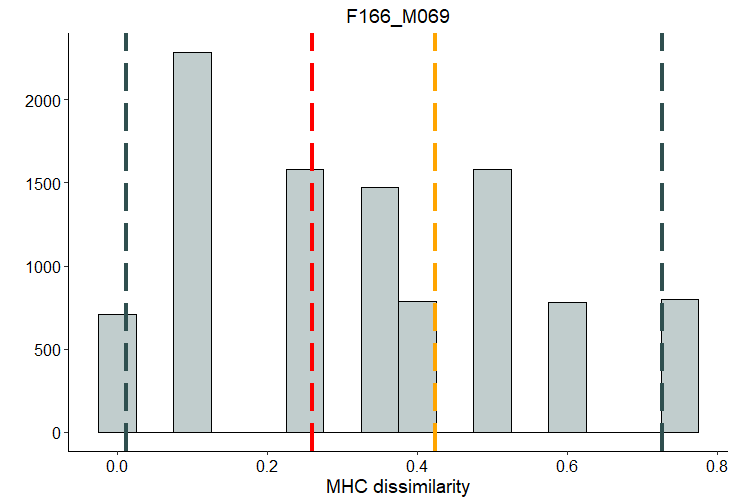  n = 13 | **c.**  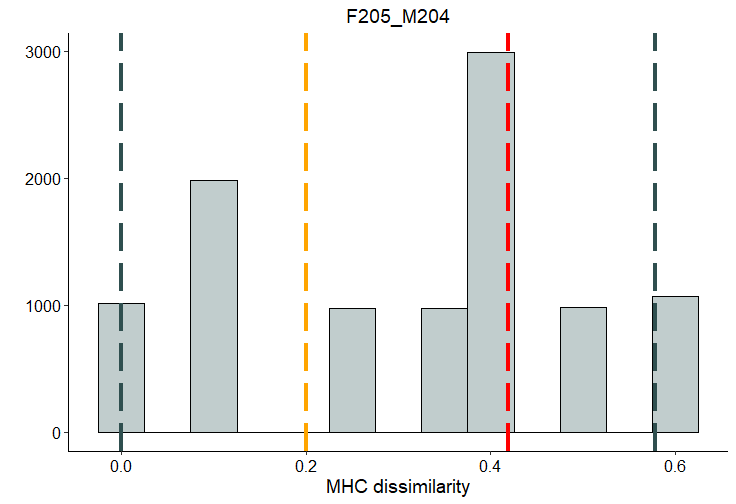 n = 10 |
| --- | --- | --- |
| **d.**  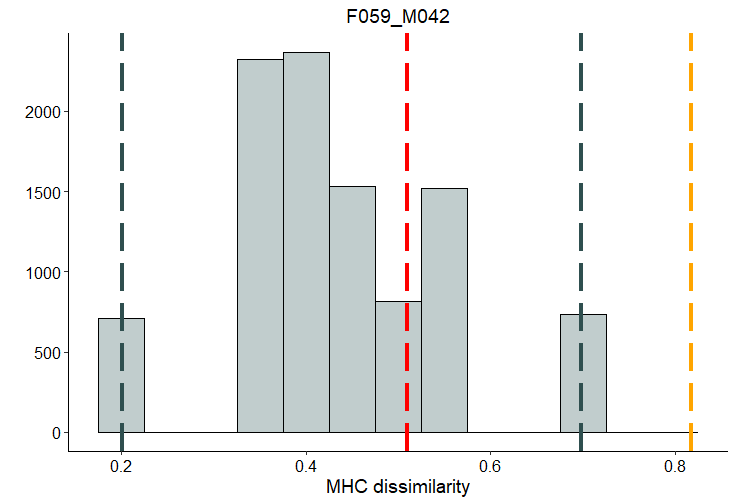 n = 13 | **e.**  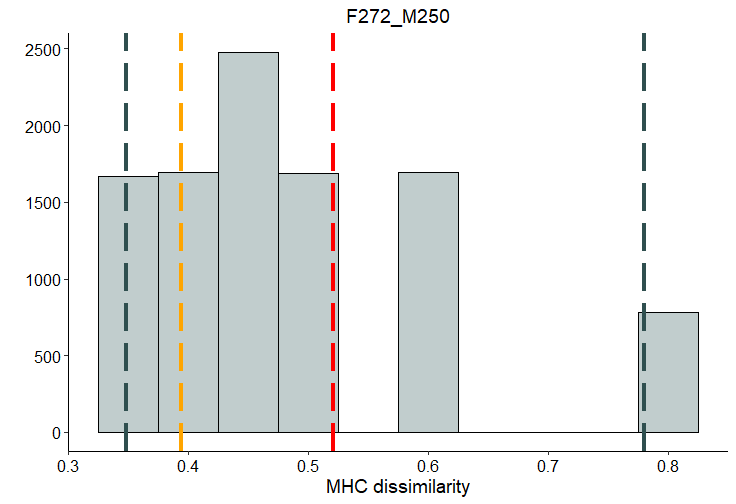 n = 12 | **f.**  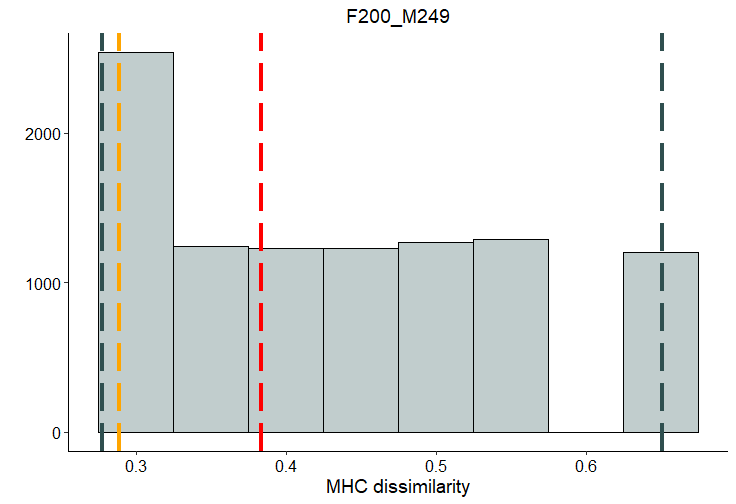 n = 18 |
| **g.**  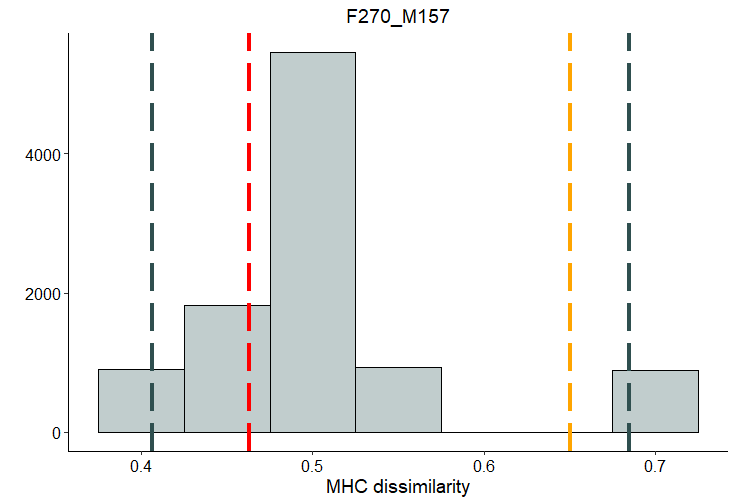  n = 11 | **h.**  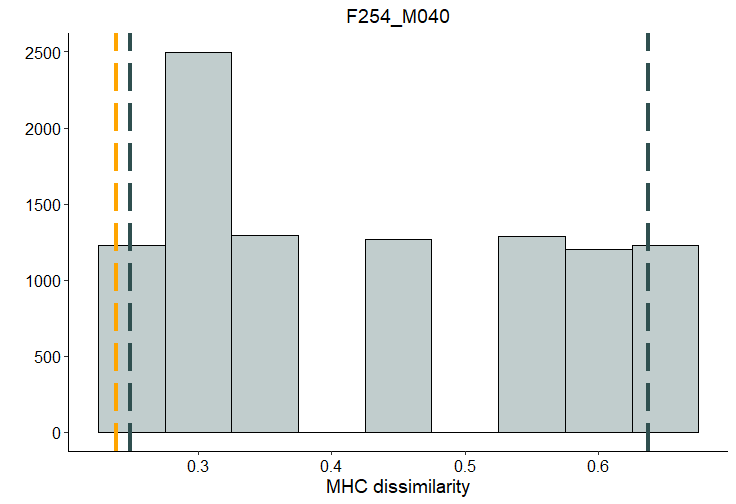  n = 8 | **i.**  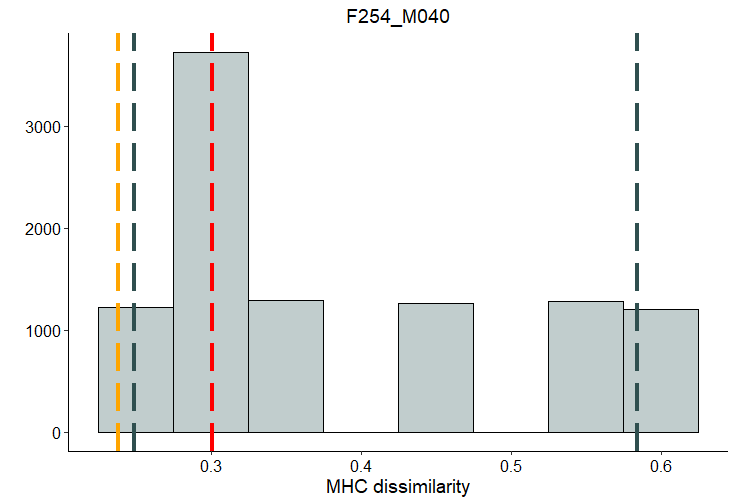  n = 8 |
| **j.**  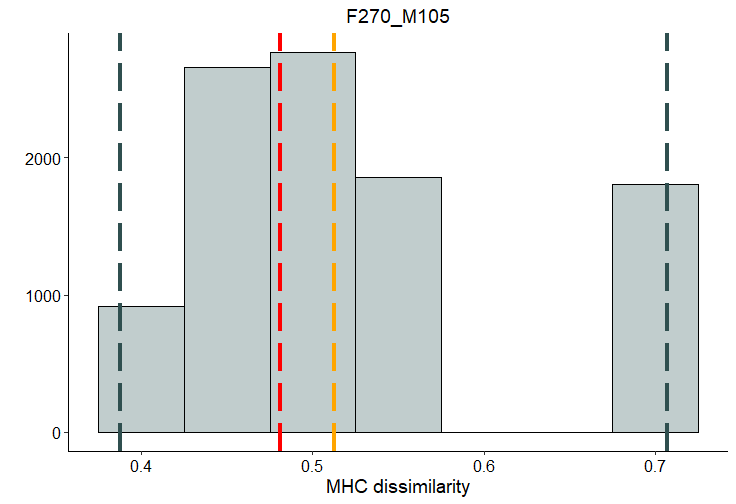  n = 11 | **k.**  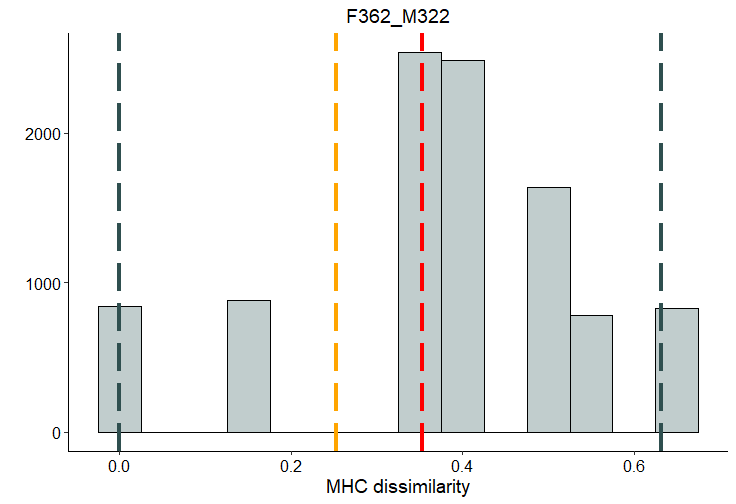  n = 12 | **l.**  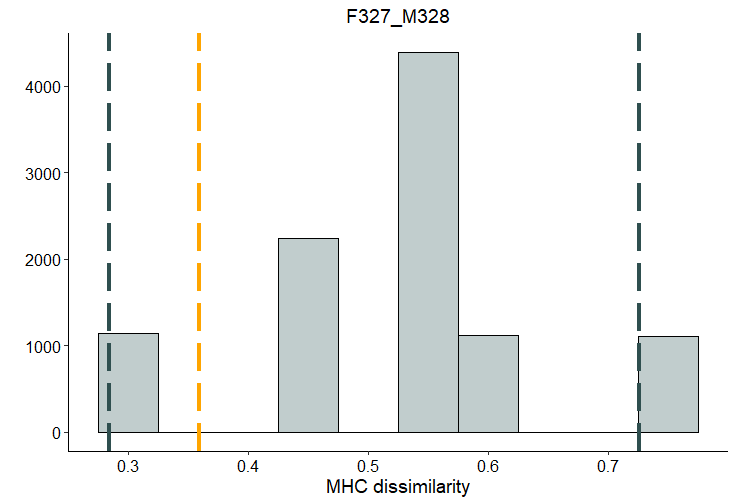  n = 9 |

| **m.**  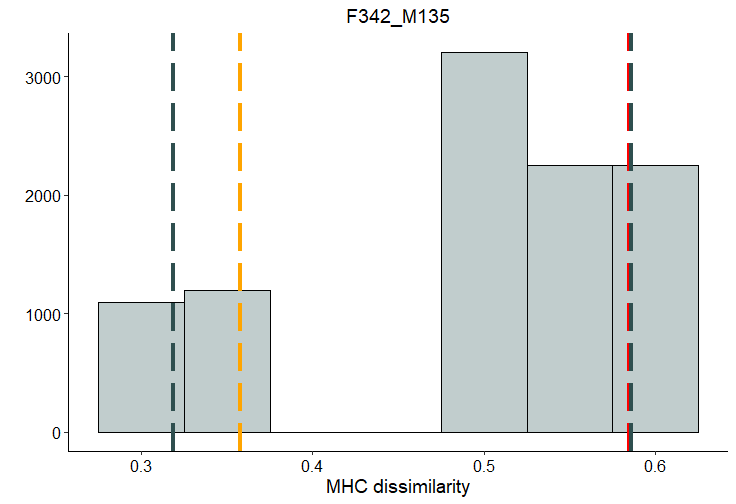  n = 9 | **n.**  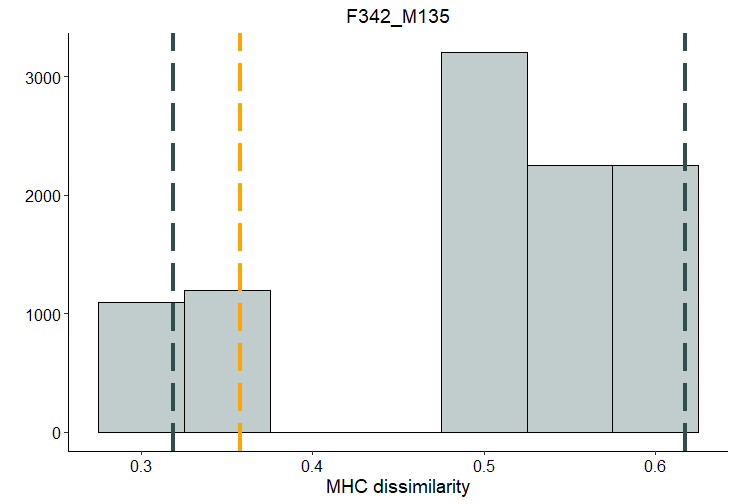  n = 9 | **o.**  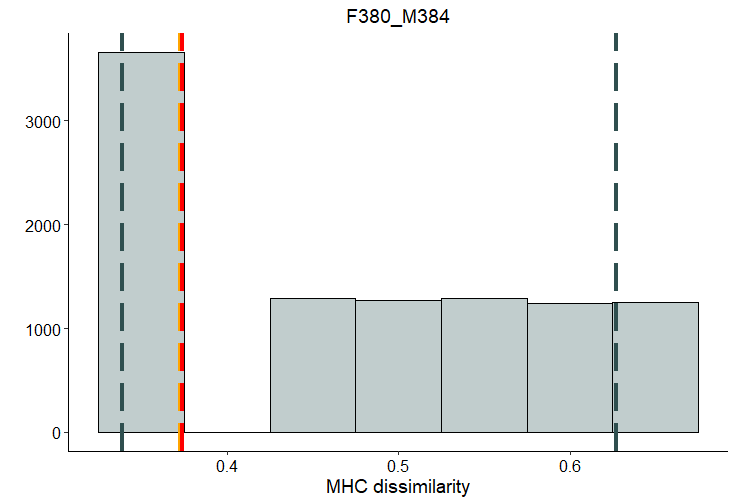  n = 8 |
| --- | --- | --- |
| **p.**  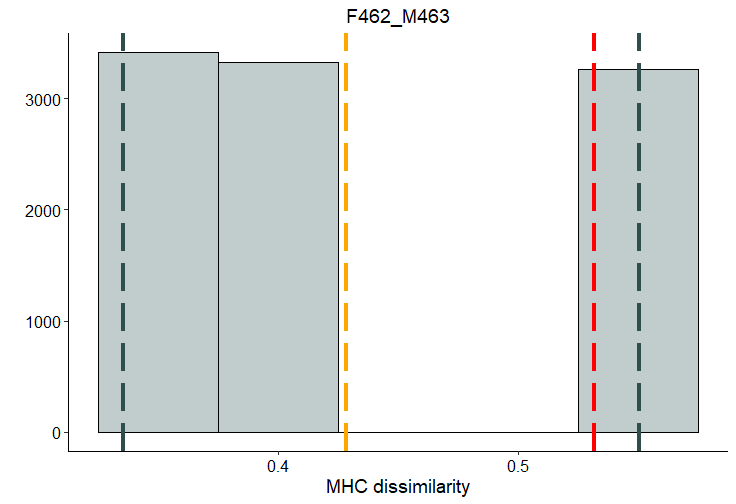  n = 6 | **q.**  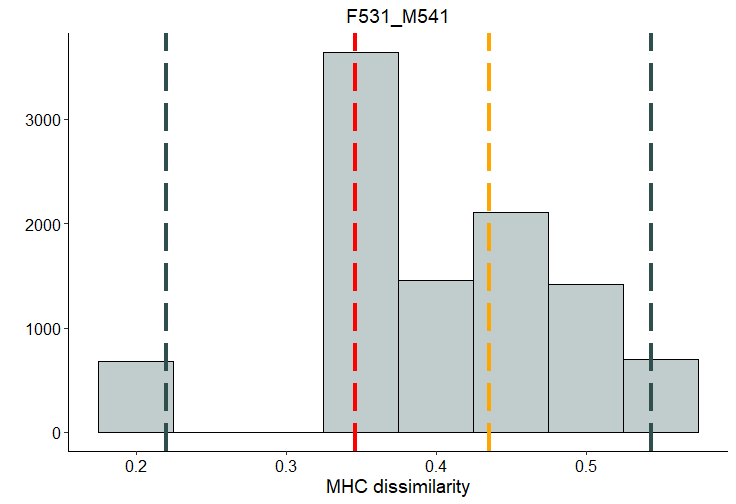  n = 14 | **r.**  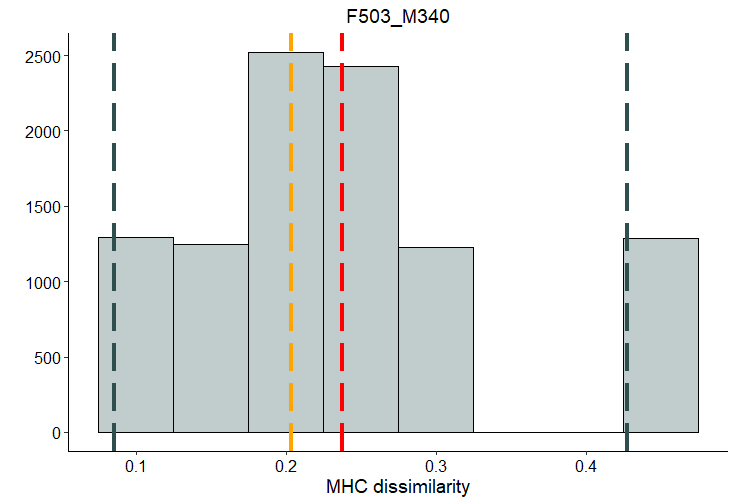  n = 8 |
| **s.**  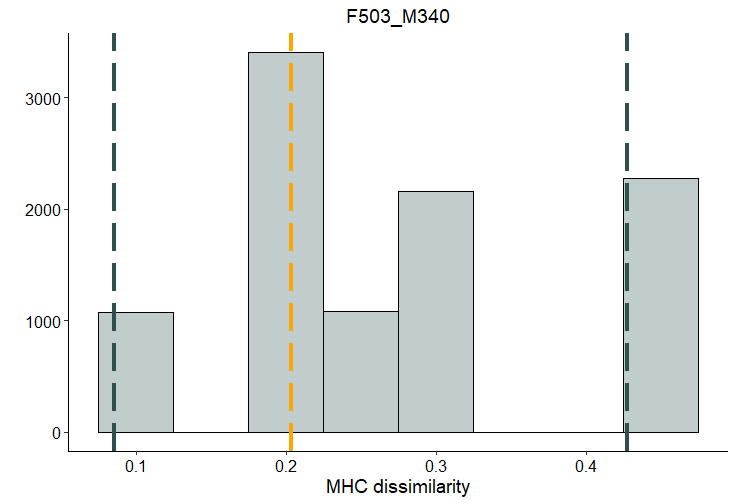  n = 9 | **t.**  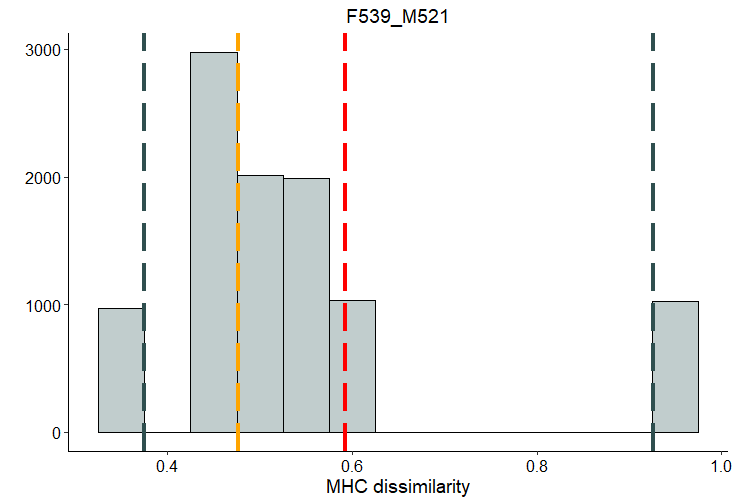  n = 10 | **u.**  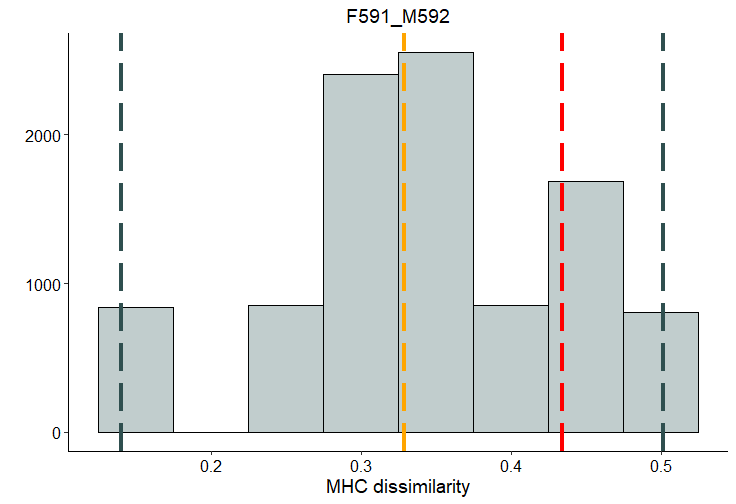  n = 12 |

**Figure S4:** Distribution of mean MHC dissimilarity scores calculated from 10000 randomly generated potential pairings between each focal female in SP-EPY and all potentially available males in the surrounding area males (6 to 14 putative males). Dark grey dashed lines depict the lower 2.5% and upper 97.5% of the data (95% confidence intervals). The mean MHC dissimilarity for the SP-EPY and EGP are depicted by the orange and red dashed lines, respectively. The order of pairs matches the order in Fig. 3 (1 to 21) in the main manuscript. *Note: when the red dashed line is not visible (as in plots h, l, n & s) the MHC dissimilarity score of the EGP is identical to either the lower (l) or upper bounds (h, n & s) of the 95% confidence intervals*.
